# G-quadruplex forming sequences in the genome of all known human viruses: a comprehensive guide

**DOI:** 10.1101/344127

**Authors:** Enrico Lavezzo, Michele Berselli, Ilaria Frasson, Rosalba Perrone, Giorgio Palù, Alessandra R. Brazzale, Sara N. Richter, Stefano Toppo

**Affiliations:** Department of Molecular Medicine, University of Padova, Padova, 35131, Italy; Department of Statistical Sciences, University of Padova, Padova, 35131, Italy

**Keywords:** G-quadruplex, viruses, secondary nucleic acid structures, bioinformatics analysis

## Abstract

G-quadruplexes are non-canonical nucleic acid structures that control transcription, replication, and recombination in organisms. G-quadruplexes are present in eukaryotes, prokaryotes, and viruses. In the latter, mounting evidence indicates their key biological activity. Since data on viruses are scattered, we here present a comprehensive analysis of putative G-quadruplexes in the genome of all known viruses that can infect humans. We show that the presence, distribution, and location of G-quadruplexes are features characteristic of each virus class and family. Our statistical analysis proves that their presence within the viral genome is orderly arranged, as indicated by the possibility to correctly assign up to two-thirds of viruses to their exact class based on the G-quadruplex classification. For each virus we provide: i) the list of all G-quadruplexes formed by GG-, GGG- and GGGG-islands present in the genome (positive and negative strands), ii) their position in the viral genome along with the known function of that region, iii) the degree of conservation among strains of each G-quadruplex in its genome context, iv) the statistical significance of G-quadruplex formation. This information is accessible from a database (http://www.medcomp.medicina.unipd.it/main_site/doku.php?id=g4virus) to allow the easy and interactive navigation of the results. The availability of these data will greatly expedite research on G-quadruplex in viruses, with the possibility to accelerate finding therapeutic opportunities to numerous and some fearsome human diseases.

## INTRODUCTION

G-quadruplexes (G4s) are nucleic acids secondary structures that may form in single-stranded DNA and RNA G-rich sequences under physiological conditions (Lipps and Rhodes 2009). Four Gs bind via Hoogsteen-type base-pairing to yield G-quartets: stacking of at least two G-quartets leads to G4 formation, through π-π interactions between aromatic systems of G-quartets. K^+^ cations in the central cavity relieve repulsion among oxygen atoms and specifically support G4 formation and stability (Sen and Gilbert 1990). In the human genome, putative DNA G4 forming regions are clustered at definite genomic regions, such as telomeres, oncogene promoters, immunoglobulin switch regions, DNA replication origins and recombination sites (Maizels and Gray 2013). Actual G4s were shown to be involved in key regulatory roles, including transcriptional regulation of gene promoters and enhancers, translation, chromatin epigenetic regulation, and DNA recombination (Perrone et al. 2013a; Holder and Hartig 2014; Maizels 2015; Rhodes and Lipps 2015; Zhou et al. 2015). Expansion of G4-forming motifs were associated with relevant human neurological disorders (Fry and Loeb 1994; Sket et al. 2015; Zhou et al. 2015). In RNA, G4s were mapped in mRNAs and in non-coding RNAs (ncRNAs), such as long non-coding RNAs (lncRNAs) (Jayaraj et al. 2012) and precursor microRNAs (pre-miRNAs) (Mirihana Arachchilage et al. 2015) indicating the potential of RNA G4s to regulate both pre- and post-transcriptional gene expression (Agarwala et al. 2015; Cammas and Millevoi 2017). RNA G4s were also suggested to affect DNA processes (Takahama et al. 2013; Hirashima and Seimiya 2015; Zheng et al. 2015).

Formation of G4s in vivo was consolidated by the discovery of cellular proteins that specifically recognize G4s (Qiu et al. 2015; Tosoni et al. 2015) and the development of G4 specific antibodies (Biffi et al. 2013; Biffi et al. 2014; Henderson et al. 2014) and compounds (Largy et al. 2013; Laguerre et al. 2015; Zheng et al. 2015; Doria et al. 2017). Experimental analyses recently attested around 10,000 G4 structures in the human genome (Hansel-Hertsch et al. 2016).

Viruses are intracellular parasites that replicate by exploiting the cell replication and protein synthesis machineries. Viruses that infect humans are very diverse and, according to the Baltimore classification, they can be divided in seven groups based on the type of their genome and mechanism of genome replication: DNA viruses with 1) double-stranded (ds) and 2) single stranded (ss) genome; RNA viruses with 3) ds genome, or ss genome with 4) positive (ssRNA (+)) or 5) negative (ssRNA (-)) polarity; 6) RNA or 7) DNA viruses with reverse transcription (RT) ability, whose genome is converted from RNA to DNA during the virus replication cycle (Table 1). Each of these classes possesses a peculiar replication cycle (Flint et al. 2015).

**Table 1.** Virus families divided according to their genome and mechanism of replication. The suffix word “viridae” for each virus family has been omitted.

| Genome nature | DNA |  | RNA |  |  | DNA and RNA |  |
| --- | --- | --- | --- | --- | --- | --- | --- |
| Group | 1 | 2 | 3 | 4 | 5 | 6 | 7 |
| Genome type | dsDNA | ssDNA | dsRNA | ssRNA (+) | ssRNA (-) | ssRNA (RT) | dsDNA (RT) |
| Virus family | Herpes | Anello | Reo | Corona | Rhabdo | Retro | Hepadna |
|  | Adeno | Parvo |  | Astro | Filo |  |  |
|  | Papilloma |  |  | Calici | Paramyxo |  |  |
|  | Polyoma |  |  | Flavi | Arena |  |  |
|  | Pox |  |  | Picorna | Bunya |  |  |
|  |  |  |  | Toga | Orthomyxo |  |  |

The presence of G4s in viruses and their involvement in virus key steps is increasingly evident (Metifiot et al. 2014; Ruggiero and Richter 2018). In the dsDNA group, the presence of G4s was described in both *Herpesviridae* and *Papillomaviridae* families. In particular, the herpes simplex virus 1 (HSV-1) possesses several repeats of important DNA G4-forming sequences (Artusi et al. 2015) that, when visualized with a G4-specific antibody in infected cells, massively form in the cell nucleus, peaking during viral replication and localizing according to the viral genome intracellular movements (Artusi et al. 2016). The HSV-1 G4s were stabilized by G4 ligands with consequent inhibition of viral DNA replication (Artusi et al. 2015; Callegaro et al. 2017). A G4 ligand and its interaction with the telomeric repeats present in the HHV-6 genome were responsible for inhibition of human herpes virus-6 (HHV-6) genome integration into the host cell chromosome (Gilbert-Girard et al. 2017). Similarly, G4 stabilizing compounds led to a reduction in Kaposi’s sarcoma associated herpesvirus (KSHV) DNA replication and disrupted maintenance of the virus latent state (Madireddy et al. 2016). In the Epstein–Barr virus (EBV), RNA G4s were implicated in the regulation of viral DNA replication and translation (Norseen et al. 2009; Murat et al. 2014) and in the delay of viral antigen-specific T cells priming (Tellam et al. 2014). In addition, the interaction of nucleolin with EBNA1 mRNA G4 mediated the virus immune evasion (Lista et al. 2017). In human papillomavirus (HPV), the presence of DNA G4s at the genome level was reported (Tluckova et al. 2013). In ssDNA viruses, the presence of G4s in the adeno-associated virus genome and their interaction with nucleophosmin were shown to repress virus entry and replication (Satkunanathan et al. 2017). In ssRNA (+) viruses, RNA G4s were described in the genomes of flaviviruses (e.g. Zika) and hepatitis C virus (HCV) (Fleming et al. 2016; Wang et al. 2016a). In the severe acute respiratory syndrome (SARS) coronavirus, a viral protein domain necessary for viral genome replication/transcription specifically binds RNA G4s (Tan et al. 2009; Kusov et al. 2015). RNA G4s were also found in the ssRNA (-) Ebola virus genome (Wang et al. 2016b). In hepatis B virus (HBV), the only member of dsDNA viruses with RT activity, stabilization of a G4 in the promoter of the envelope gene increased transcription and virion secretion (Biswas et al. 2017). Functionally significant G4s were identified in the human immunodeficiency virus (HIV), a retrovirus belonging to group 6 (Table 1). G4s were described both in the RNA and DNA proviral genome (Perrone et al. 2013a; Perrone et al. 2014; Piekna-Przybylska et al. 2014): in the latter, clusters of G4s were reported in the unique long terminal repeat (LTR) promoter (Perrone et al. 2013a; Amrane et al. 2014) and in the Nef coding region (Perrone et al. 2013b). The cellular proteins nucleolin and hnRNP A2/B1 regulate the G4/ds equilibrium in the LTR promoter (Tosoni et al. 2015; Lago et al. 2017; Scalabrin et al. 2017) and G4 ligands induced significant antiviral effects in a G4-mediated mechanism of action (Perrone et al. 2013a; Perrone et al. 2014; Perrone et al. 2015). The LTR region of lentiviruses in general (ssRNA RT) present stable and conserved G4s (Perrone et al. 2017a).

Given this amount of scattered data, we here aimed at analyzing the presence of putative G4 forming patterns in the genome of all known viruses that can cause infections in humans. The analysis is performed at two distinct levels, globally for each viral genome and individually for each detected G4 pattern. First, we asked the following: is the number of putative G4s found in a viral genome simply due to chance, hence trivially reflecting genomic G/C content? Are putative G4s scattered on the genome or preferentially found in regulatory regions? To address these questions, we collected the whole viral genomes deposited in databanks, we clustered them in multiple sequence alignments, and searched for the presence of G4s looking for their conservation in variable/constant genomic regions. The results were compared with an *in silico* generated set of simulated genomes containing the same nucleotide composition of the corresponding real ones. Results show that many viruses have a highly significant content of putative G4s and their distribution is not random but rather peculiar to specific genomic regions (e.g. coding sequences, repeats, regulatory regions). Once attested that the presence of putative G4s is significant for some viruses, we further analyzed every single identified G4 in a virus species for its conservation across virus strains, as shown in Figure 1 reporting the data analysis workflow. The detailed information on the putative G4s present in each human virus is available in an easily accessible web site with interactive graphics for the straightforward interpretation and navigation of the data (http://www.medcomp.medicina.unipd.it/main_site/doku.php?id=g4virus).

**Figure 1.**
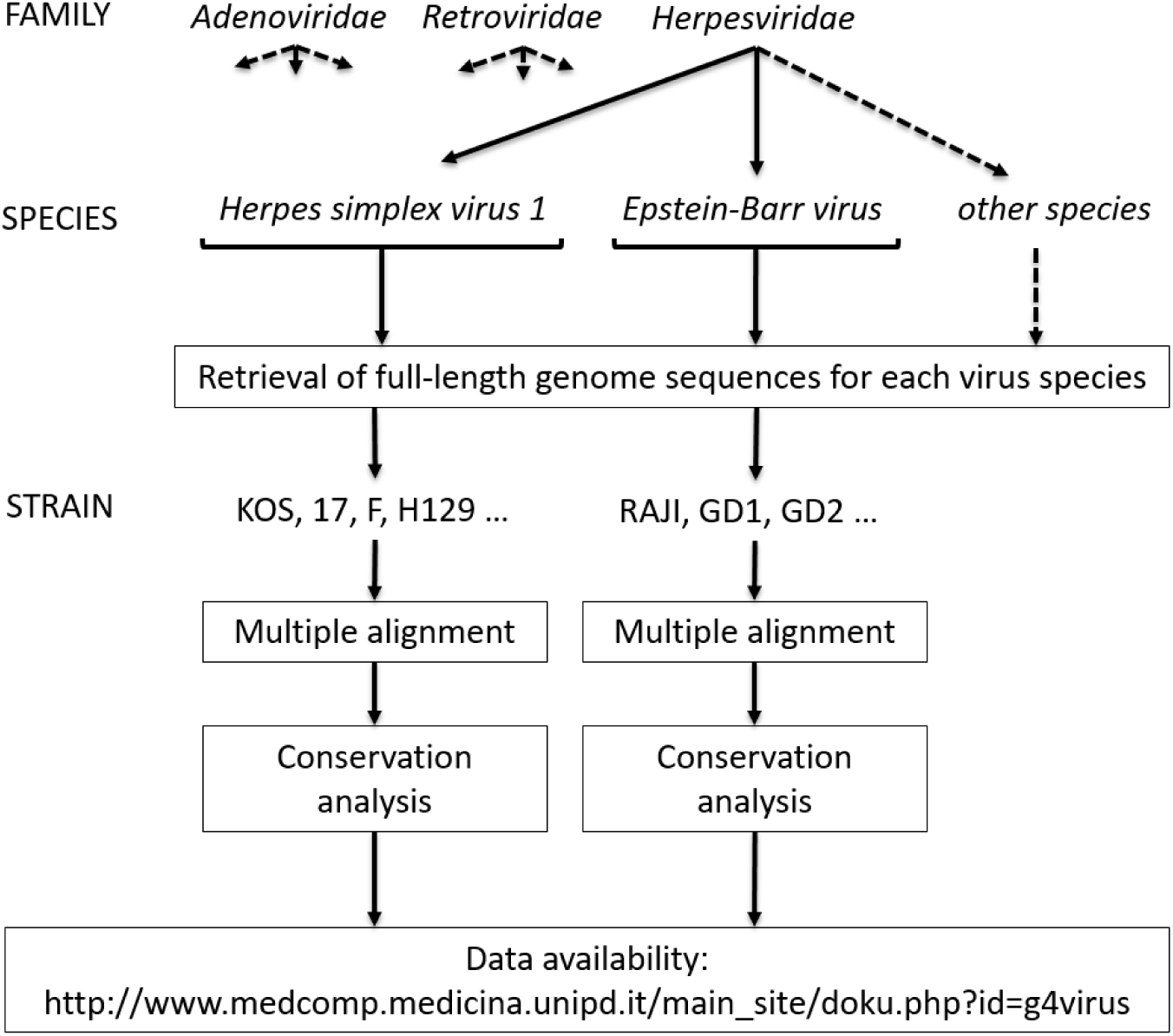
Example of virus classification in Families, Species and Strains with data analysis flowchart. The conservation analysis was performed among the strains within each virus species.

In addition, we observed that the G4s content can also be a distinctive feature of some viral species hence can be exploited for their classification in the corresponding Baltimore group. To investigate this aspect, we used a semiparametric classifier to assign the virus to its exact class relying on its G4 content and we observed that G4-assignment is statistically relevant in most viruses. G4 features are characteristic of virus classes and families, to such an extent that up to two-thirds of the viruses could be correctly classified into their virus class based on the G4 features.

To our knowledge, this is the first comprehensive study that describes the presence and distribution of G4s in human viruses. In our idea, this work will serve as a guidance for scientists interested in studying the role of G4s in viruses, providing a valid instrument for the selection of the most interesting candidates for experimental validations.

## RESULTS

### Detection of G4 patterns in all known human viruses

All known viruses that cause infections in humans, according to the Viral Zone ExPASy web site (http://viralzone.expasy.org/all_by_species/678.html), were grouped in 7 classes according to Baltimore classification, which takes into account the viral genome nature: dsDNA, ssDNA, dsRNA, ssRNA(+), ssRNA(-), ssRNA(RT) and dsDNA(RT). Different replication strategies and structural similarities allow to further divide viruses in families (Table 1). The complete list of reference sequences for each virus included in the analyses is reported in Supplemental_Material.pdf (Table S1).

G4 patterns were searched in the positive and negative strand of each virus genome sequence, since both filaments are present and important in different stages of the viral replicative cycle of all virus classes. As the length of virus genomes greatly varies, i.e. from 235,646 nt of the human cytomegalovirus (HCMV) to 1,682 nt of hepatitis delta virus (HDV), we reported the number of putative G4 sequences independently of the genome length by normalizing their number per 1,000 nts (Figure 2). The G4 pattern distribution for both the positive and negative strands is shown as a box plot for each Baltimore virus class, whereas the G4 count for each virus within each class is shown as a dot besides the box plot (Figure 2). The negative strand of retroviruses (ssRNA (RT) viruses), ssDNA viruses and both strands of dsDNA viruses showed the largest presence of G4s made of GG-, GGG- and GGGG-islands (box plots, Figure 2). Both strands of genomes of single virus families belonging to these groups and to ssRNA (+) and ssRNA (-) were enriched in G4s of all G-islands types (dot plots, Figure 2). Conversely, dsRNA and dsDNA (RT) viruses notably lacked the presence of G4s.

**Figure 2.**
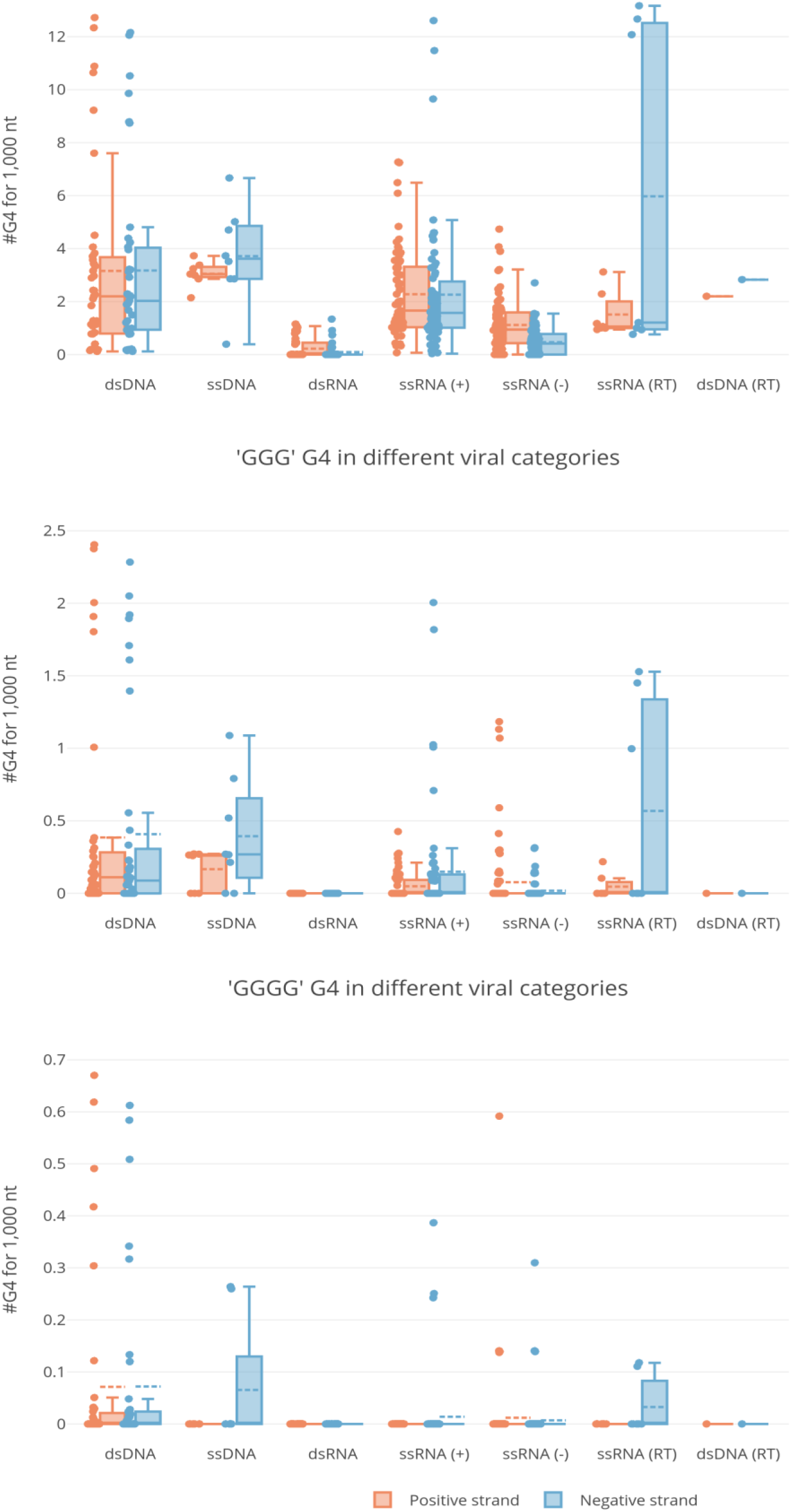
Box and whisker plots with outliers of G4s in different virus classes. Each panel refers to the indicated type of G-island (GG, GGG, GGGG). The abundance of G4s per 1 kb of viral genome is reported in the y-axis and the different virus categories in the x-axis. Boxplots are delimited by the first and third quartile and the straight and dotted lines drawn inside are the median and mean values, respectively, of the G4 distribution. The single observations are reported as dots close to the box plot. Whiskers delimit all the points that fall above/below the third/first quartile plus/minus 1.5 times the interquartile range (IQR). Orange and blue box plots refer to positive and negative strand respectively.

We next reasoned about the conservation of G4 patterns among different strains of each viral species, hypothesizing that the presence of a conserved G4 pattern within a less conserved genome environment could be an indication of a G4 with a biological function. To allow for the evaluation of G4 conservation in the local context of viral genomes, we computed the “G4 scaffold conservation index” (G4_SCI) for each G4 in each virus species. This value measures the degree of conservation of G-islands that are necessary and sufficient to form a G4: the higher the score, the higher the conservation of the G4 pattern. An example of the results from such analysis is reported in Figure 3 for both the lymphocytic choriomeningitis virus (segment S) and the human coxsackievirus A: all G4s for each virus are plotted as vertical bars, the height and position of which represent the G4_SCI on the y-axis and the genome coordinates on the x-axis, respectively. In addition, the local sequence conservation (LSC) of the viral genome, calculated with a sliding window approach on all available viral sequences and plotted as a red broken line, is reported alongside. This visualization method allows the prompt identification of the presence, position, and conservation of G-islands within G4s, together with the overall local conservation of the genomic context. Moreover, the degree of conservation of the connecting regions (loops) with respect to G-islands (the *loop_conservation* value) was calculated as the difference between G4_SCI and LSC. Positive and negative *loop_conservation* scores indicate, respectively, lower and higher conservation of connecting regions compared to the conservation of G-islands. Values close to zero mean that both G-islands and connecting loops show the same level of sequence conservation. In Figure 3, three G4s formed by highly conserved GG-islands are shown for the S segment of lymphocytic choriomeningitis virus, present in genomic regions both well and less well conserved (Figure 3 at positions 1,790 in the positive strand, 1,759 and 2,676 in the negative strand). Similarly, the human coxsackievirus A genome displays four G4s formed by GG-islands that are extremely conserved and embedded in a less conserved genomic environment (i.e. positions 3,725, 5,891, 5,834 in the positive strand, 2,926 in the negative strand) and one highly conserved G4 in a highly conserved region (i.e. position 458 in the negative strand) (Figure 3). This kind of analysis is available for all G4s of all human virus species at http://www.medcomp.medicina.unipd.it/mainsite/doku.php?id=g4virus (*loop_conservation* values are included in tarballs downloadable for each viral class of the Baltimore classification, whereas each virus species has a dedicated page displaying all graphical representations).

**Figure 3.**
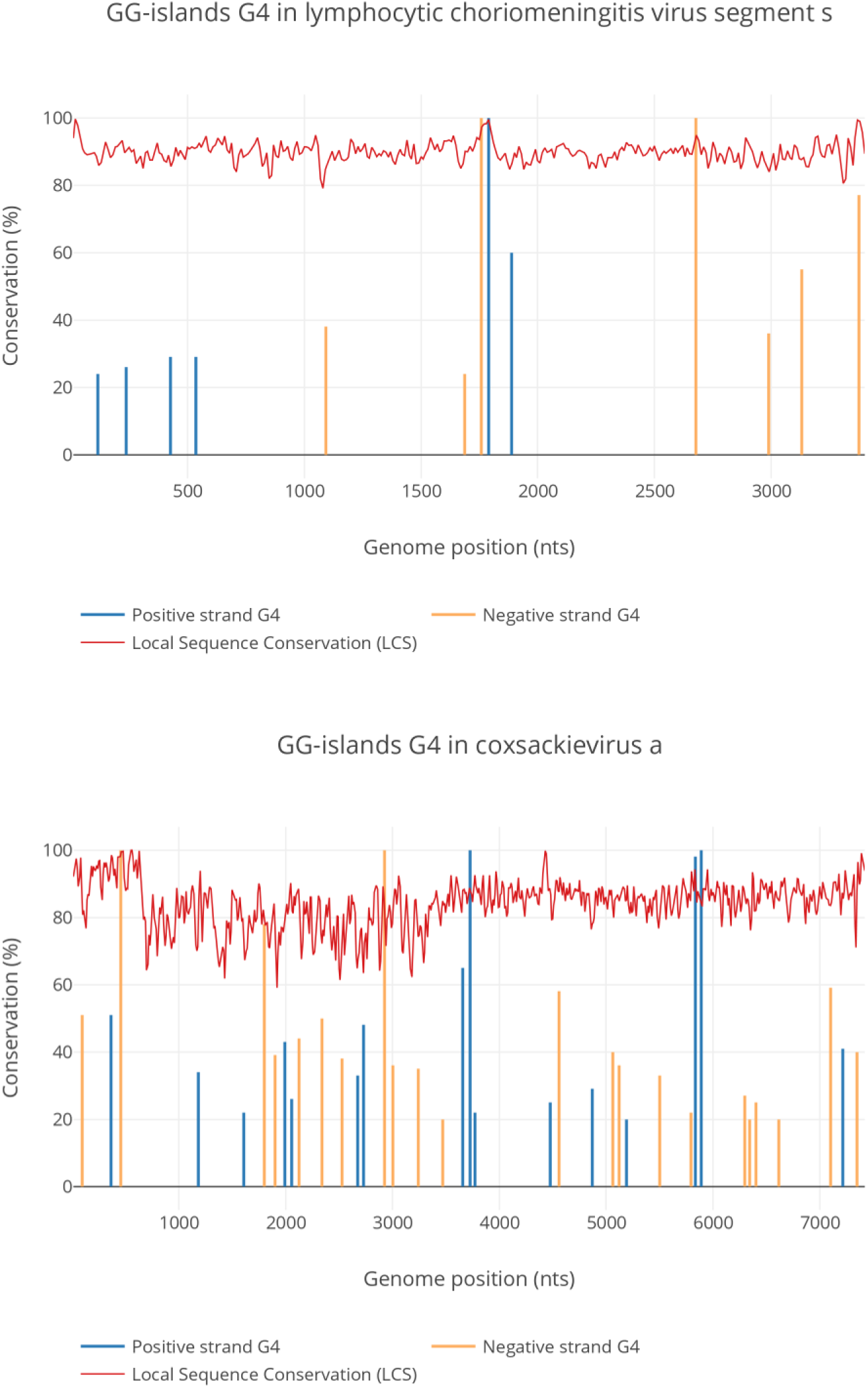
Conservation of G4s and viral genomes. G4s formed by GG-islands in the S segment of lymphocytic choriomeningitis virus and in coxsackievirus A are shown. G4s found in the positive and negative strands are indicated as blue and orange vertical bars, respectively, while the height of bars represents the G4_SCI (the conservation of G-islands). Local sequence conservation (LSC) of viral genomes is shown as a red broken line. The x-axis indicates genome position, the y-axis the conservation %. This information is available for all viruses at http://www.medcomp.medicina.unipd.it/mainsite/doku.php?id=g4virus.

To assess the results, we retrieved from the literature all the available experimentally validated G4s detected in human viruses. All patterns were confirmed also by our analysis and the complete list is reported in Supplemental_Material.pdf (Table S2), together with the genomic coordinates of the predicted G4s.

### Statistical evidence for the calculated G4s in the human virus genomes

G4 formation may be largely affected by G/C content, which greatly varies in viral genomes (from 76% of Cercopithecine 2 herpes virus to 27% of Yaba like disease virus). Moreover, it has been shown that some di- and trinucleotides are over or underrepresented in certain viruses (Greenbaum et al. 2008; Lobo et al. 2009) and, in the context of G4 patterns, it means that their abundance could be biased by unexpected frequencies of guanine homopolymers. G-island frequencies higher or lower than expected would lead to a potential over- or under-representation of G4s, respectively.

To check whether the presence of putative G4s was statistically relevant or whether it occurred by pure chance, we compared the results obtained from real viral genomes with those obtained by two different simulation strategies. The first one (single nucleotide assembling) assumes that the occurrence of each DNA base in the genome is independent (Huppert and Balasubramanian 2005); the second (G-island reshuffling) considers that short sequences of a given length (k-mer) could be over or underrepresented in certain viral genomes (Greenbaum et al. 2008; Lobo et al. 2009). In the former case, sequences were generated with the same composition of nucleotides but different order with respect to references; in the latter, sequences were produced by reshuffling the positions of G-islands while keeping constant their number.

For each virus and simulation strategy, we produced 10,000 random sequences, which were screened with our G4s detection pipeline. Real and simulated data were compared by computing a P-value, defined as twice the smaller proportion of simulated sequences that exhibit, respectively, a higher and lower count of G4s as compared to the median value of all the available complete genome sequences for a certain virus. Hence, a P-value close to 1 means that the median G4 content in real viral sequences is not significant if compared to a random distribution; conversely, a P-value close to 0 means that G4 content is highly significant. This interpretation holds independently of the length of the genome and/or of the prevalence of either G/C bases or G-islands, as we compare the number of G4s in a viral genome with the one we would expect in a simulated genome of the same length and of either the same base or G-island composition. To account for possible high discreteness of the data, a less conservative version of the P-value, called the mid-P value (Armitage and Berry 1994), was used. Segment diagrams of the mid-P values of the Baltimore grouped viruses are reported in Supplemental_Material.pdf (Figure S1a and Figure S1b) (Chambers et al. 1983). The number of viruses whose median G4 count is significant at the 10% level is listed in Table 2 (virus names in Supplemental_Table_S3.xlsx) with the indication of whether this median count is either higher or lower than the G4 count in simulated sequences.

**Table 2.** Relative abundance of viruses having a G4 pattern content significantly different between real and simulated viral genomes. The number of viruses where the amount of G4 patterns is significantly different at 10% level between real and simulated sequences is reported (with percentages in brackets). Values and percentages were calculated considering only viruses containing at least one G4 pattern either in real or simulated sequences. Table reports significant values for either one of the two simulations (randomization of viral genomes at single nucleotide or at G-island levels) or both.

|  | Number of viruses significantly different vs. randomization at single nucleotide level |  | Number of viruses significantly different vs. randomization at the G-island level |  | Number of viruses significantly different vs. both randomization |  |
| --- | --- | --- | --- | --- | --- | --- |
| G-island pattern | G4 more abundant in real sequences | G4 more abundant in simulated sequences | G4 more abundant in real sequences | G4 more abundant in simulated sequences | G4 more abundant in real sequences | G4 more abundant in simulated sequences |
| **GG (positive strand)** | 83/218 (38.1%) | 9/218 (4.1%) | 52/217 (24.0%) | 4/217 (1.8%) | 49/217 (22.6%) | 3/217 (1.4%) |
| **GG (negative strand)** | 83/187 (44.4%) | 2/187 (1.1%) | 67/187 (35.8%) | 1/187 (0.5%) | 59/186 (31.7%) | 0 (0%) |
| **GGG (positive strand)** | 41/78 (52.6%) | 3/78 (3.8%) | 40/75 (53.3%) | 0 (0%) | 34/74 (45.9%) | 0 (0%) |
| **GGG (negative strand)** | 32/68 (47.1%) | 3/68 (4.4%) | 32/69 (46.4%) | 0 (0%) | 28/68 (41.2%) | 0 (0%) |
| **GGGG**<br>**(positive**<br>**strand)** | 17/19<br>(89.5%) | 0 (0%) | 17/17<br>(100%) | 0 (0%) | 17/17<br>(100%) | 0 (0%) |
| **GGGG**<br>**(negative**<br>**strand)** | 23/28<br>(82.1%) | 0 (0%) | 23/25<br>(92.0%) | 0 (0%) | 23/25<br>(92.0%) | 0 (0%) |

Our data show that most members of the dsDNA, ssDNA, and ssRNA (RT) present a highly significant content of G4s formed by GG-, GGG- and/or GGGG-islands in one or both strands. ssRNA (-) and ssRNA (+) classes are heterogeneous since some viruses are highly significant in any G4 category (from GG- to GGGG-islands), while others are not (see below). Formation of G4s in members of the dsRNA group is notably less significant. Interestingly, few viruses display a smaller amount of G4 patterns than expected: both *Sagiyama virus* and *Human coronavirus HKU1* are depleted of G4s belonging to GG-islands category in the positive genome strand when compared with both simulation strategies based on single nucleotide assembling and GG-island reshuffling. In addition, *Human parainfluenza virus 2* is poor of G4 patterns made of GG-island in the positive genome strand but is enriched in both GG- and GGG-type G4s in the negative strand.

Overall, if we consider the viruses that contain at least one G4 pattern in either the real or the simulated genomes, we observe that the increase in G4 islands’ length corresponds to a decrease in the absolute number of viruses containing G4s, but it also corresponds to a dramatic increase in the fraction of them that is statistically significant.

By looking at the family level of viral classification, which is far more homogeneous than the Baltimore groups, some virus families emerge as prominently enriched in G4 patterns. Among them, *Herpesviridae* is not only the one with the highest G4 content, but most of its members display significantly more G4 patterns than expected in both genome strands and in all considered G-island lengths. Notably, some of the viruses belonging to *Herpesviridae* and showing the highest G/C content are statistically enriched in G4s. This suggests that simply having a high G/C content is not a sufficient condition to justify the presence of such a high number of G4s. Other viral families that are consistently enriched in G4s are *Adenoviridae* and *Papillomaviridae*, especially in GG- (both strands) and GGG-island (positive strand) types. *Poxviridae* and *Parvoviridae* show an enrichment of GG-type G4s in both genome strands, whereas the same pattern is enriched in the positive strand of all *Anelloviridae* members and in the negative strand of most *Paramyxoviridae* and *Retroviridae* viruses. All other families are generally not enriched in G4s in any of the evaluated categories, with only a few exceptions that are listed in the following: L segments of *Lassa virus* and *Lymphocytic choriomeningitis virus* (*Arenaviridae*), *Wu* and *Merkel cell polyomaviruses* (*Polyomaviridae*), *Salivirus* (*Picornaviridae*), M and S segments of respectively *Crimean-Congo hemorrhagic fever virus* and *Rift Valley fever virus* (*Bunyaviridae*).

By comparing the results obtained independently from the two simulation strategies it is possible to draw additional conclusions. First, in most cases the results are concordant, meaning that both simulations show similar trends in the statistical significance. Nonetheless, the overall number of viruses whose G4 content is significantly different with respect to simulated data is higher when real viral genomes are compared to those generated by single nucleotide assembling. This difference indicates that viral genome k-mer composition is indeed affecting the probability of randomly finding G4 patterns, at least in a proportion of viruses as shown in Figure 4: in the heatmaps, viruses that are significant in only one of the two simulations are reported for GG- and GGG-island patterns, whereas no such cases were found for GGGG-type G4s.

**Figure 4.**
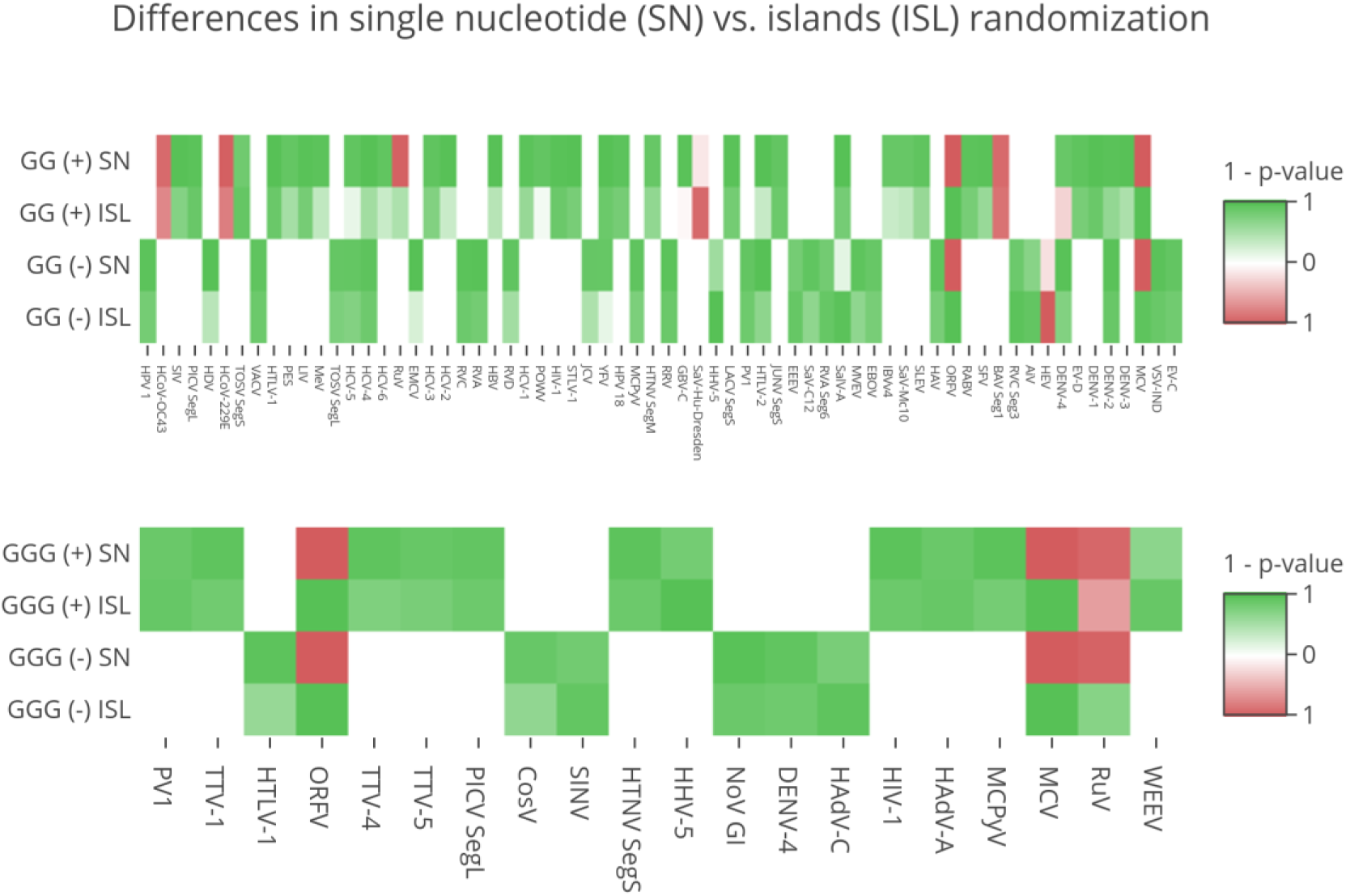
Different results from single nucleotide (SN) and islands (ISL) reshuffling strategies. Heatmaps show all the viruses which are significant in only one of the two simulations or that obtain discordant results. Green and red boxes indicate that G4s are more abundant in real and simulated genomes, respectively, with color intensity proportional to p-value size.

Finally, some remarkable exceptions exist where both simulations return a significant p-value, but with an opposite meaning. This is the case of two members of the *Poxviridae* family, namely *Molluscum contagiosum virus* and *Orf virus*, which are enriched in GG- and GGG-type G4s in both strands of their genomes if compared with the islands reshuffling simulation but show the opposite behavior when compared with the single nucleotide assembling (they are also reported in Figure 4). While the full meaning of this observation is not clear to us, it seems that these viruses possess far less G4 patterns than they could have, but at the same time they are able to cluster their relatively few G-islands in more G4 patterns than expected.

### Human virus G4 patterns position and overlap with genomic features

To check the prevalent positions of G4s in virus genomes, we compared the coordinates of predicted G4s with the available information regarding viral genome features. Genome coordinates were extracted for coding sequences (CDS), repeat regions (RR), 5’- and 3’-untranslated (UTR), and promoter regions. While CDS and RR are explicitly defined in RefSeq and GenBank databases, the annotation of UTRs and promoters is more inconsistent, being defined only for some viral species. For this reason, the annotations of genes and CDSs were exploited to indirectly extract the coordinates of 5’– and 3’–regulatory regions (see Materials and Methods for details). To determine the localization of G4s in viral genomes, the overlap extent between G4 patterns and genomic features was computed. Given the vast heterogeneity of the annotations reported in the feature fields, a manual revision was required to fix potential inconsistencies in annotations, regarding both keywords and coordinates. A revision was performed when possible, while controversial and uncertain annotations were not considered. These analyses are presented as bar charts for individual viral classes and G4-island pattern types (GG-, GGG-, GGGG-island) (Supplemental_Material.pdf, Figure S2a-c). As regards the GGG-island type, the herpesvirus family of dsDNA viruses presents G4s distributed along all the four identified genomic features, with a particularly high concentration in RR and, in some members, in the 5’ – regulatory region. This feature is consistent with the reported extent of G4s in HSV-1, which is mainly clustered in the RR of the virus genome (Artusi et al. 2015; Artusi et al. 2016). Conversely, viruses belonging to the ssRNA (+) and ssRNA (-) classes show G4s mainly grouped in CDS and in the 3’- and 5’-regulatory regions, respectively. HIV-1, belonging to ssRNA (RT) virus class, presents G4s of the GGG-island type mainly in the RR and 3’-regulatory regions and in part in CDS. This distribution confirms previous data (Perrone et al. 2013a; Perrone et al. 2013b). Conversely, other retroviruses (ssRNA (RT)) such as HTLV-1 and HTLV-2, display G4s in the CDS. Given the lower stringency of G4s of the GG-island type, these are more widely distributed along the four identified genomic features, whereas the most stringent G4s of the GGGG type, present only in herpesviruses (dsDNA) and HTLV-1 (ssRNA (RT)), show a clear-cut localization in the RR and CDS, respectively. These data indicate that the distribution of putative G4s in the viral genomes is not random and their localization differs in virus classes.

### The number and type of putative G4s are characteristic of virus classes

In this line of thinking, we asked whether the observed number of G4s, and more precisely its statistical significance with respect to the two random assembling scenarios, is representative for a particular virus class. To answer this question, we checked whether it is possible to classify each virus of the six classes considered, that is, dsDNA, ssDNA, dsRNA, ssRNA (+), ssRNA (-) and ssRNA (RT), into its own class based on the information of how significant its median G4 counts are. We chose to use a classifier built on multinomial logistic regression, as this method is both interpretable and robust to unbalanced group sizes as long as the group sizes are large enough. To avoid the latter drawback, we excluded from the model fit the hepatitis B virus, the only virus classified as dsDNA (RT), and the two unclassified Hepatitis delta and Hepatitis E viruses. Six features were used to classify the viruses, i.e. the six mid-P values (those calculated for GG-, GGG-, GGGG-, both in the positive and negative strand) which qualify the G4 content of the real viral sequences. The values were multiplied by 1 or −1 depending on whether the median G4 count was over- or underrepresented. Since real and corresponding simulated sequences contain the same base or G-islands composition, the classification model based on G4 content does not depend on the highly variable content of G/C in the different virus classes but is specifically designed on the peculiar presence or absence of G4s in each viral class. Furthermore, 34 viruses with no G4 count in all three G-island types in both the positive and negative strand and non-significant mid-P values at the 10% level were excluded from the analysis. We reclassified every viral genome used in our assessment using the discriminant function obtained from a leave-one-out analysis. This latter technique allowed us to accurately estimate how our classifier performs without the need to split our data into a training and a test set. The corresponding confusion matrix is given in Table 3 from where it is possible to extract the overall percentage of correct classifications that amount to 66.7% for the single nucleotide assembling model and 68.1% for the G-island reshuffling model. The agreement is good for the dsDNA, ssDNA, dsRNA, ssRNA (+) and ssRNA (-) classes. The two unclassified genomes of the Hepatitis delta and Hepatitis E viruses were classified as ssRNA (+).

**Table 3.** Confusion matrix for the semi-parametric classifier for G4 structure. The classifier is based on a multinomial model and uses the one-sided mid-P values as features in combination with the information on whether the median G4 count is under- or over-represented.

| Single nucleotide assembling model |  | Predicted class |  |  |  |  |  |  |
| --- | --- | --- | --- | --- | --- | --- | --- | --- |
|  |  | dsDNA | ssDNA | dsRNA | ssRNA (+) | ssRNA (-) | ssRNA (RT) | Unclassified |
| True class | dsDNA | 33 | 0 | 0 | 1 | 4 | 1 | 0 |
|  | ssDNA | 2 | 0 | 0 | 5 | 0 | 1 | 0 |
|  | dsRNA | 0 | 0 | 13 | 0 | 5 | 0 | 0 |
|  | ssRNA (+) | 4 | 0 | 0 | 45 | 13 | 0 | 0 |
|  | ssRNA (-) | 3 | 0 | 8 | 18 | 48 | 0 | 0 |
|  | ssRNA (RT) | 4 | 0 | 0 | 0 | 0 | 3 | 0 |
|  | Unclassified | 0 | 0 | 0 | 2 | 0 | 0 | 0 |
| G-island reshuffling model |  | Predicted class |  |  |  |  |  |  |
|  |  | dsDNA | ssDNA | dsRNA | ssRNA (+) | ssRNA (-) | ssRNA (RT) | Unclassified |
| True class | dsDNA | 31 | 0 | 0 | 5 | 1 | 2 | 0 |
|  | ssDNA | 4 | 0 | 0 | 2 | 2 | 0 | 0 |
|  | dsRNA | 0 | 0 | 14 | 0 | 4 | 0 | 0 |
|  | ssRNA (+) | 3 | 1 | 0 | 48 | 10 | 0 | 0 |
|  | ssRNA (-) | 7 | 1 | 4 | 13 | 52 | 0 | 0 |
|  | ssRNA (RT) | 3 | 0 | 0 | 3 | 1 | 0 | 0 |
|  | Unclassified | 0 | 0 | 0 | 2 | 0 | 0 | 0 |

## DISCUSSION

In this work we provide: i) the list of all G4s present in all human virus genomes (both positive and negative strands), ii) their position in the viral genome along with the known function of that part of the genome (i.e. coding, regulating, repeated regions), iii) the degree of conservation of both G4-islands and loops vs. the genome, iv) the statistical significance of G4 formation in each virus.

Our data show that viruses belonging to dsDNA, ssDNA, ssRNA (RT) and, to a less extent, ssRNA (+) and ssRNA (-) display the largest presence of GG-, GGG- and GGGG-type G4s (box plots, Figure 2) and that the presence of G4s in all these virus groups is statistically significant (segment diagrams, Supplemental_Material.pdf, Figure S1a and Figure S1b). Both results support a role of G4s in the virus biology: indeed, some G4s predicted in this work were already reported in viruses and were shown to possess specific and different functions.

We evidenced some general trends and exceptions that are worth noting if seen in comparative terms among all viruses considered in this study. Starting from the general features we noted: i) high G/C content is not sufficient per se to generate a high number of G4s, as observed in G/C rich members of *Herpesviridae* family that are richer of putative G4s than expected. ii) The statistical significance of putative G4s found in real sequences tends to decrease when G-islands reshuffling (ISL) is compared with the corresponding single nucleotide assembling (SN), as is appreciable from the heatmap in Figure 4 (more intense color in the heatmap boxes). This suggests that short sequences of a given length (k-mer) could be over or underrepresented in certain viral genomes, as already reported in the literature (Greenbaum et al. 2008; Lobo et al. 2009). We observed that viral genomes enriched in putative G4s also contain a higher number of G-islands than expected from mere nucleotide composition, especially evident in the GG-islands. iii) The unevenly distribution of G4s can be used to classify membership of a virus in its corresponding category. This was not predictable *a priori* but up to two-thirds of unequivocal assignments suggest that for some viruses the G4 content works as a distinctive feature. iv) G4 localization shows differences in some virus classes but this outcome is still incomplete due to lack of information in the databases about virus genomic features and partitioning into regulatory and coding regions.

Some other interesting observations are worth reporting as either special cases or exceptions. To start with, the ssRNA (-) group is the most heterogeneous one, since some viral species are significantly enriched in G4s up to the most extended G-island type (GGGG), while others lack this feature. Surprisingly, two viruses of the dsDNA group, which was generally highly enriched in G4s, show a significantly lower presence of putative G4 forming patterns than expected in a random sequence with the same G/C content (SN, Supplemental_Table_S3.xlsx), even though the opposite result was observed vs. simulated genomes with the same G-islands content as the real ones (ISL). These two viruses, i.e. *Molluscum contagiosum virus* and *Orf virus*, are the only ones belonging to genera other than the *Orthopoxvirus* within the *Poxviridae* family that cause skin lesions. Finally, dsRNA and dsDNA (RT) viruses are notably poor in G4s and with mostly null statistical significance; however, single G4s are highly conserved (e.g. rotavirus a segment 6), therefore still conveying potential biological interest.

These data indicate that G4s are mainly present in ss-genome viruses, which in principle are more suitable to fold into G4s since they do not require unfolding from a fully complementary strand. The major exception to this evidence is the *Herpesviridae* family of dsDNA viruses. In this case, most G4s reported here and also previously described (Artusi et al. 2015; Artusi et al. 2016) form in repeated regions. It is possible that repeated sequences are more prone to alternative folding, as shown by several non-canonical structures that form in repeated region of the DNA (Fry and Loeb 1994; Reddy et al. 2013; Sket et al. 2015; Zhou et al. 2015). However, for some herpesviruses a large number of G4s is also present in regulatory regions, which may indicate yet undiscovered functional roles. To note that the investigation of putative G4 forming sequences was performed on a maximum window of 48 nucleotides: however it is possible, especially for the ss genomes, that bases more distant in the primary sequence interact among each other, therefore expanding the repertoire of G4 structures.

The significant enrichment of G4s in many viruses with respect to the corresponding randomized genomes is an indication that the clustering of G-islands did not occur by pure chance, suggesting a specific biological role of the G4 structures. Complementary to this, the analysis of the G4 conservation highlights every G4 that is conserved among viral strains. Since one of the peculiarities of viral genomes is their fast mutation rate, the strong conservation of a specific G-island pattern among strains is per se an indication of the biological relevance of a G4 forming sequence. In light of this, single conserved G4s in viruses that do not display statistically significant G4 enrichment may nonetheless retain key functional roles.

The discrepancy between the conservation of G-islands and connecting loops (*loop_conservation*) is an additional indication on the likeliness of biological implications of a specific G4. A positive *loop_conservation* value highlights G4-islands more conserved than their connecting loops, suggesting that only the G4 scaffold is required for mechanisms that are important for the virus life cycle, while the loops are dispensable. Considering the high mutation rate of viruses, this type of conservation indicates sequences where G4 formation is most likely essential. Equally conserved G4-islands and loops (*loop_conservation* value = 0) imply that both the G4 scaffold and connecting loops are potentially relevant for the virus and probably involve interactors that specifically recognize them. In this case, the high sequence conservation, especially in CDS, may depend on the required conservation of that peculiar gene product rather than the presence of a G4 structure. Nonetheless, the option of targeting these conserved G4s for therapeutic purposes remains unaltered and the availability of specific and conserved loops may only enhance the chance of finding selective ligands. Therefore, from this point of view, the ‘zero’ class is the best scenario for the development of specific drugs. The “negative” *loop_conservation* value scenario is of less immediate interpretation: it is possible that false positive hits fall in this category as it is unexpected that G4-islands are less conserved than their connecting loops.

The evidence provided here, the previous studies on G4s in viruses, and the possibility to correctly classify the majority of viruses based on their G4 features (Table 3) suggest that most of the virus classes adopted G4-mediated mechanisms to control their viral cycles.

This work offers comprehensive data to guide researchers in the choice of the most significant G4s within a human virus genome of interest. Hopefully this will accelerate research in this area with the identification of new G4-mediated mechanisms in viruses and the development of effective and innovative therapeutics.

## MATERIAL AND METHODS

The complete list of viral species able to infect humans was retrieved from http://viralzone.expasy.org/allbyspecies/678.html (accessed in April 2016) and, for each of them, all available complete genome sequences were downloaded from GenBank. Multiple alignments were built for each virus with usearch8 (Edgar 2010), using a permissive identity threshold (60%) to account for viral variability. Since in some cases nucleotide heterogeneity within viral species exceeded this value, multiple clusters of aligned sequences were obtained for some viruses, representing distinct genotypes. Considering the difficulty of obtaining high quality alignments beyond this limit of nucleotide similarity, all clusters obtained with this method were kept separate, manually assigned to specific genotypes and independently processed in the downstream analyses. One genome per each group of aligned sequences was chosen to serve as reference sequence, possibly belonging to the manually curated RefSeq database [https://www.ncbi.nlm.nih.gov/refseq/]. The complete list of selected reference sequences is reported in Supplemental_Material.pdf (Table S1).

G4 patterns were searched in all nucleotide sequences with an in-house developed tool, as previously described (Perrone et al. 2017a; Perrone et al. 2017b). Briefly, a G4 was reported when at least four consecutive guanine islands (G-islands) were detected. If ‘*l’* is the minimum number of G in every G-island of a G4 and ‘*d’* is the maximum distance allowed between two consecutive G-islands, the following combinations of ‘*l’* and ‘*d’* were searched: *l* = 2 and *d* = 7; *l* = 3 and *d* = 12; *l* = 4 and *d* = 12. Patterns in the negative strands of viral genomes were searched by looking for cytosines (Cs) instead of Gs. The conservation of each G4 in the multiple aligned genomes of the viruses was determined by looking at the conservation not only of the G-islands but also their connecting loops. We computed different indexes to measure the nucleotide sequence conservation of viral genomes and G4s:

*i) G4_scaffold_conservation_index* (*G4_SCI*): it is an index referred to the G-islands. For each virus and for every detected G4 pattern, it is calculated as the percentage of independent genomes maintaining the corresponding G-islands:

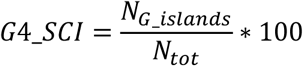

where N_G_islands_ is the number of sequences possessing the G-islands in a certain genome position and N_tot_ is the total number of sequences available for the virus.

*ii) Loop_conservation*: it is the difference between G4_SCI and the local conservation of the viral sequence spanning the G4 (LSC_G4_).

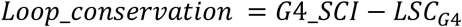

LSC_G4_ is calculated as the average of LSC windows overlapping the G4. LSC measure is computed within a sliding window of fixed length (length 20, shift 10), averaging the conservation values of each position extracted from the multiple sequence alignments with Jalview (Waterhouse et al. 2009). They are formally defined as:

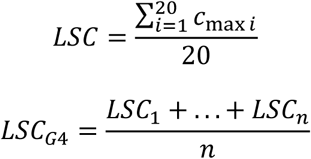

where *c_max i_* is the maximum conservation at position *i* of the multiple aligned sequences and *n* is the number of windows overlapping the G4. The results of these analyses are presented individually for each virus and G4 pattern (http://www.medcomp.medicina.unipd.it/main_site/doku.php?id=g4virus), together with the calculated profiles of simple linguistic complexity and Shannon entropy that can highlight other potential local features of the sequence (e.g. repeats and low complexity regions) (Berselli et al. 2018). All charts were generated with Plotly [https://plot.ly], exploiting Pandas (McKinney 2010) and Numpy Python libraries (van der Walt et al. 2011). Unless otherwise stated, analyses were conducted with ad hoc developed Python and Perl scripts, which are available in the website (scripts.tar.gz).

### Evaluation of G4 conservation in real vs randomized viral sequences

To determine whether the presence of G4 patterns in a virus is a conserved feature or it is only a consequence of its nucleotide composition, simulated viral genomes were generated and compared with real data. Two different strategies were adopted to generate simulated data:

i) *Single nucleotide assembling (SN).* A computational approach was adopted where, in analogy to Huppert and Balasubramanian (Huppert and Balasubramanian 2005), the viral genome was modelled as a multinomial stream based on the assumption that each DNA base is independent. These authors give an explicit solution for the prevalence of G4s in the human genome as a function of *p(G)*, the probability of any base being G. In our approach, we also accounted for the probability of cytosines (*p(C)*) and additionally assumed that adenine (A) and thymine (T) bases were equally likely to occur. As all four probabilities need to sum up to one, the statistical reference model is a multinomial distribution with probability vector *(p(G), p(C), p(A), p(T))*. We hence took as many independent draws from this multinomial distribution as the number of nucleotides in the reference viral genome (Supplemental_Material.pdf, Table S1). The probabilities *p(G)* and *p(C)* vary for each virus and reflect the prevalence of G and C bases present in that virus, while the remaining proportion is equally split to give *p(A)* and *p(T)*. For each virus, 10,000 independent sequences were produced *in silico* with this method; the ‘sample’ R command with replacement was used and the provided parameters were the genome length and the relative abundance of the four nucleotides in the real genomes.

ii) *G-islands reshuffling (ISL).* For each virus, we generated a scaffold of length *n* made of only As, with *n* corresponding to the length of the virus reference genome; then, we replaced di-, tri-, or tetra-nucleotides, at random positions and without overlaps, with as many GG-, GGG-, GGGG-, CC-, CCC-, CCCC-islands as in the reference sequence, respectively. Overall, we generated 10,000 independent sequences for three different simulated datasets, one for each island length.

### Virus classification assignment by G4 content

The simulated sequences were scanned for the presence of G4 patterns as previously described. The 10,000 counts obtained for each simulation formed the empirical distribution for G4 prevalence under the hypothesis of random assembling of the genome in the SN and ISL models respectively. The mid-P value was calculated using a homemade function. The semiparametric classifier used to assign the virus to its exact class relying on its G4 content was based on the ‘multinom’ function of the R package ‘nnet’.

### G4 patterns position and overlap with genomic features

The feature tables containing viral genome annotations were downloaded from RefSeq or GenBank for all the reference sequences reported in Supplemental_Material.pdf (Table S1). Genome coordinates were extracted for coding sequences (‘CDS’), repeat regions (‘repeat_region’), 5’- and 3’-untranslated (UTR) and promoter regions. Given the annotation inconsistency of promoters and UTRs, two new feature categories were created, 5’ – and 3’ – regulatory regions that were defined by exploiting the annotation of genes and CDSs. We calculated boundaries for genes in the positive strand of viral genomes as follows: 5’ – regulatory = S_gene_ – S_cds_; 3’ – regulatory = E_cds_ - E_gene_. For the genes in the negative strand of viral genomes we defined: 5’ – regulatory = S_cds_ - S_gene_; 3’ – regulatory = E_gene_ - E_cds_. S_gene_, S_cds_, E_gene_ and E_cds_ are the Start (S) and End (E) of genes and CDSs as extracted from the feature tables. These newly defined features contain both UTRs and promoters. Since the positive genomic strands are deposited in RefSeq for most of the viruses belonging to the ssRNA (-) class, the following sequences available as negative strands were converted into their inverse complement, together with the coordinates of their genomic features: Junin arenavirus segment S (NC_005081) and segment L (NC_005080), Lassa virus segment L (NC_004297), lymphocytic choriomeningitis virus segment S (GQ862982), Machupo virus segment S (AY924208) and L (AY624354), Pichinde virus segment S (NC_006447), Rift Valley fever virus segment S (NC_014395), and Toscana virus segment S (NC_006318). The overlap extent between G4 patterns and genomic features was computed by intersecting the genomic coordinates of each G4 pattern with the genomic features extracted from the corresponding virus, and a positive count was recorded every time an overlap of at least one nucleotide was detected. Finally, to compare the enrichment in different feature classes, characterized by different sizes, a normalization step was performed. The total extension of each feature class (*i.e.* CDS, repeat_region, 5’ – regulatory and 3’ – regulatory) was calculated by summing the lengths of individual features. The total count of G4 overlapping a feature class was then divided by the total length of the class and multiplied by a factor 1,000 to obtain the number of G4 present every 1,000 nucleotides. This procedure was performed considering only the G4 patterns conserved in at least 80% of sequences for each viral species. All feature tables files were manually revised to fix inconsistencies in the use of keywords and coordinates.

## ACKNOWLEDGEMENTS

This work was supported by the Bill and Melinda Gates Foundation [GCE grant numbers OPP1035881, OPP1097238]; European Research Council [ERC Consolidator grant 615879] to S.N.R. University of Padova grant CPDA138081/13 to S.T.

## REFERENCES

Agarwala P, Pandey S, Maiti S. 2015. The tale of RNA G-quadruplex. Organic & biomolecular chemistry 13(20): 5570–5585.

Amrane S, Kerkour A, Bedrat A, Vialet B, Andreola ML, Mergny JL. 2014. Topology of a DNA G-quadruplex structure formed in the HIV-1 promoter: a potential target for anti-HIV drug development. Journal of the American Chemical Society 136(14): 5249–5252.

Armitage P, Berry G. 1994. Statistical Methods in Medical Research.

Artusi S, Nadai M, Perrone R, Biasolo MA, Palu G, Flamand L, Calistri A, Richter SN. 2015. The Herpes Simplex Virus-1 genome contains multiple clusters of repeated G-quadruplex: Implications for the antiviral activity of a G-quadruplex ligand. Antiviral research 118: 123–131.

Artusi S, Perrone R, Lago S, Raffa P, Di Iorio E, Palu G, Richter SN. 2016. Visualization of DNA G-quadruplexes in herpes simplex virus 1-infected cells. Nucleic acids research 44(21): 10343–10353.

Berselli M, Lavezzo E, Toppo S. 2018. NeSSie: a tool for the identification of approximate DNA sequence symmetries. Bioinformatics.

Biffi G, Di Antonio M, Tannahill D, Balasubramanian S. 2014. Visualization and selective chemical targeting of RNA G-quadruplex structures in the cytoplasm of human cells. Nature chemistry 6(1): 75–80.

Biffi G, Tannahill D, McCafferty J, Balasubramanian S. 2013. Quantitative visualization of DNA G-quadruplex structures in human cells. Nature chemistry 5(3): 182–186.

Biswas B, Kandpal M, Vivekanandan P. 2017. A G-quadruplex motif in an envelope gene promoter regulates transcription and virion secretion in HBV genotype B. Nucleic acids research.

Callegaro S, Perrone R, Scalabrin M, Doria F, Palu G, Richter SN. 2017. A core extended naphtalene diimide G-quadruplex ligand potently inhibits herpes simplex virus 1 replication. Scientific reports 7(1): 2341.

Cammas A, Millevoi S. 2017. RNA G-quadruplexes: emerging mechanisms in disease. Nucleic acids research 45(4): 1584–1595.

Chambers J, Cleveland W, Kleiner B, Tukey P. 1983. Graphical Methods for Data Analysis.

Doria F, Nadai M, Zuffo M, Perrone R, Freccero M, Richter SN. 2017. A red-NIR fluorescent dye detecting nuclear DNA G-quadruplexes: in vitro analysis and cell imaging. Chemical communications 53(14): 2268–2271.

Edgar RC. 2010. Search and clustering orders of magnitude faster than BLAST. Bioinformatics 26(19): 2460–2461.

Fleming AM, Ding Y, Alenko A, Burrows CJ. 2016. Zika Virus Genomic RNA Possesses Conserved G-Quadruplexes Characteristic of the Flaviviridae Family. ACS infectious diseases 2(10): 674–681.

Flint SJ, Racaniello VR, Glenn FR, Skalka AM, Enquist LW. 2015. Principles of Virology: Volume 1 Molecular Biology. ASM Press.

Fry M, Loeb LA. 1994. The fragile X syndrome d(CGG)n nucleotide repeats form a stable tetrahelical structure. Proceedings of the National Academy of Sciences of the United States of America 91(11): 4950–4954.

Gilbert-Girard S, Gravel A, Artusi S, Richter SN, Wallaschek N, Kaufer BB, Flamand L. 2017. Stabilization of Telomere G-Quadruplexes Interferes with Human Herpesvirus 6A Chromosomal Integration. Journal of virology 91(14).

Greenbaum BD, Levine AJ, Bhanot G, Rabadan R. 2008. Patterns of evolution and host gene mimicry in influenza and other RNA viruses. PLoS pathogens 4(6): e1000079.

Hansel-Hertsch R, Beraldi D, Lensing SV, Marsico G, Zyner K, Parry A, Di Antonio M, Pike J, Kimura H, Narita M et al. 2016. G-quadruplex structures mark human regulatory chromatin. Nature genetics 48(10): 1267–1272.

Henderson A, Wu Y, Huang YC, Chavez EA, Platt J, Johnson FB, Brosh RM, Jr., Sen D, Lansdorp PM. 2014. Detection of G-quadruplex DNA in mammalian cells. Nucleic acids research 42(2): 860–869.

Hirashima K, Seimiya H. 2015. Telomeric repeat-containing RNA/G-quadruplex-forming sequences cause genome-wide alteration of gene expression in human cancer cells in vivo. Nucleic acids research 43(4): 2022–2032.

Holder IT, Hartig JS. 2014. A matter of location: influence of G-quadruplexes on Escherichia coli gene expression. Chemistry & biology 21(11): 1511–1521.

Huppert JL, Balasubramanian S. 2005. Prevalence of quadruplexes in the human genome. Nucleic acids research 33(9): 2908–2916.

Jayaraj GG, Pandey S, Scaria V, Maiti S. 2012. Potential G-quadruplexes in the human long non-coding transcriptome. RNA biology 9(1): 81–86.

Kusov Y, Tan J, Alvarez E, Enjuanes L, Hilgenfeld R. 2015. A G-quadruplex-binding macrodomain within the “SARS-unique domain” is essential for the activity of the SARS-coronavirus replication-transcription complex. Virology 484: 313–322.

Lago S, Tosoni E, Nadai M, Palumbo M, Richter SN. 2017. The cellular protein nucleolin preferentially binds long-looped G-quadruplex nucleic acids. Biochimica et biophysica acta 1861(5 Pt B): 1371–1381.

Laguerre A, Hukezalie K, Winckler P, Katranji F, Chanteloup G, Pirrotta M, Perrier-Cornet JM, Wong JM, Monchaud D. 2015. Visualization of RNA-Quadruplexes in Live Cells. Journal of the American Chemical Society 137(26): 8521–8525.

Largy E, Granzhan A, Hamon F, Verga D, Teulade-Fichou MP. 2013. Visualizing the quadruplex: from fluorescent ligands to light-up probes. Topics in current chemistry 330: 111–177.

Lipps HJ, Rhodes D. 2009. G-quadruplex structures: in vivo evidence and function. Trends in cell biology 19(8): 414–422.

Lista MJ, Martins RP, Billant O, Contesse MA, Findakly S, Pochard P, Daskalogianni C, Beauvineau C, Guetta C, Jamin C et al. 2017. Nucleolin directly mediates Epstein-Barr virus immune evasion through binding to G-quadruplexes of EBNA1 mRNA. Nature communications 8: 16043.

Lobo FP, Mota BE, Pena SD, Azevedo V, Macedo AM, Tauch A, Machado CR, Franco GR. 2009. Virus-host coevolution: common patterns of nucleotide motif usage in Flaviviridae and their hosts. PloS one 4(7): e6282.

Madireddy A, Purushothaman P, Loosbroock CP, Robertson ES, Schildkraut CL, Verma SC. 2016. G-quadruplex-interacting compounds alter latent DNA replication and episomal persistence of KSHV. Nucleic acids research 44(8): 3675–3694.

Maizels N. 2015. G4-associated human diseases. EMBO reports 16(8): 910–922.

Maizels N, Gray LT. 2013. The G4 genome. PLoS genetics 9(4): e1003468.

McKinney W. 2010. Data Structures for Statistical Computing in Python. Proceedings of the 9th Python in Science Conference: 51–56.

Metifiot M, Amrane S, Litvak S, Andreola ML. 2014. G-quadruplexes in viruses: function and potential therapeutic applications. Nucleic acids research 42(20): 12352–12366.

Mirihana Arachchilage G, Dassanayake AC, Basu S. 2015. A potassium ion-dependent RNA structural switch regulates human pre-miRNA 92b maturation. Chemistry & biology 22(2): 262–272.

Murat P, Zhong J, Lekieffre L, Cowieson NP, Clancy JL, Preiss T, Balasubramanian S, Khanna R, Tellam J. 2014. G-quadruplexes regulate Epstein-Barr virus-encoded nuclear antigen 1 mRNA translation. Nature chemical biology 10(5): 358–364.

Norseen J, Johnson FB, Lieberman PM. 2009. Role for G-quadruplex RNA binding by Epstein-Barr virus nuclear antigen 1 in DNA replication and metaphase chromosome attachment. Journal of virology 83(20): 10336–10346.

Perrone R, Butovskaya E, Daelemans D, Palu G, Pannecouque C, Richter SN. 2014. Anti-HIV-1 activity of the G-quadruplex ligand BRACO-19. The Journal of antimicrobial chemotherapy 69(12): 3248–3258.

Perrone R, Doria F, Butovskaya E, Frasson I, Botti S, Scalabrin M, Lago S, Grande V, Nadai M, Freccero M et al. 2015. Synthesis, Binding and Antiviral Properties of Potent Core-Extended Naphthalene Diimides Targeting the HIV-1 Long Terminal Repeat Promoter G-Quadruplexes. Journal of medicinal chemistry 58(24): 9639–9652.

Perrone R, Lavezzo E, Palu G, Richter SN. 2017a. Conserved presence of G-quadruplex forming sequences in the Long Terminal Repeat Promoter of Lentiviruses. Scientific reports 7(1): 2018.

Perrone R, Lavezzo E, Riello E, Manganelli R, Palu G, Toppo S, Provvedi R, Richter SN. 2017b. Mapping and characterization of G-quadruplexes in Mycobacterium tuberculosis gene promoter regions. Scientific reports 7(1): 5743.

Perrone R, Nadai M, Frasson I, Poe JA, Butovskaya E, Smithgall TE, Palumbo M, Palu G, Richter SN. 2013a. A dynamic G-quadruplex region regulates the HIV-1 long terminal repeat promoter. Journal of medicinal chemistry 56(16): 6521–6530.

Perrone R, Nadai M, Poe JA, Frasson I, Palumbo M, Palu G, Smithgall TE, Richter SN. 2013b. Formation of a unique cluster of G-quadruplex structures in the HIV-1 Nef coding region: implications for antiviral activity. PloS one 8(8): e73121.

Piekna-Przybylska D, Sullivan MA, Sharma G, Bambara RA. 2014. U3 region in the HIV-1 genome adopts a G-quadruplex structure in its RNA and DNA sequence. Biochemistry 53(16): 2581–2593.

Qiu J, Wang M, Zhang Y, Zeng P, Ou TM, Tan JH, Huang SL, An LK, Wang H, Gu LQ et al. 2015. Biological Function and Medicinal Research Significance of G-Quadruplex Interactive Proteins. Current topics in medicinal chemistry 15(19): 1971–1987.

Reddy K, Zamiri B, Stanley SY, Macgregor RB, Jr., Pearson CE. 2013. The disease-associated r(GGGGCC)n repeat from the C9orf72 gene forms tract length-dependent uni- and multimolecular RNA G-quadruplex structures. The Journal of biological chemistry 288(14): 9860–9866.

Rhodes D, Lipps HJ. 2015. G-quadruplexes and their regulatory roles in biology. Nucleic acids research 43(18): 8627–8637.

Ruggiero E, Richter SN. 2018. G-quadruplexes and G-quadruplex ligands: targets and tools in antiviral therapy. Nucleic acids research.

Satkunanathan S, Thorpe R, Zhao Y. 2017. The function of DNA binding protein nucleophosmin in AAV replication. Virology 510: 46–54.

Scalabrin M, Frasson I, Ruggiero E, Perrone R, Tosoni E, Lago S, Tassinari M, Palu G, Richter SN. 2017. The cellular protein hnRNP A2/B1 enhances HIV-1 transcription by unfolding LTR promoter G-quadruplexes. Scientific reports 7: 45244.

Sen D, Gilbert W. 1990. A sodium-potassium switch in the formation of four-stranded G4-DNA. Nature 344(6265): 410–414.

Sket P, Pohleven J, Kovanda A, Stalekar M, Zupunski V, Zalar M, Plavec J, Rogelj B. 2015. Characterization of DNA G-quadruplex species forming from C9ORF72 G4C2-expanded repeats associated with amyotrophic lateral sclerosis and frontotemporal lobar degeneration. Neurobiology of aging 36(2): 1091–1096.

Takahama K, Takada A, Tada S, Shimizu M, Sayama K, Kurokawa R, Oyoshi T. 2013. Regulation of telomere length by G-quadruplex telomere DNA- and TERRA-binding protein TLS/FUS. Chemistry & biology 20(3): 341–350.

Tan J, Vonrhein C, Smart OS, Bricogne G, Bollati M, Kusov Y, Hansen G, Mesters JR, Schmidt CL, Hilgenfeld R. 2009. The SARS-unique domain (SUD) of SARS coronavirus contains two macrodomains that bind G-quadruplexes. PLoS pathogens 5(5): e1000428.

Tellam JT, Zhong J, Lekieffre L, Bhat P, Martinez M, Croft NP, Kaplan W, Tellam RL, Khanna R. 2014. mRNA Structural constraints on EBNA1 synthesis impact on in vivo antigen presentation and early priming of CD8+ T cells. PLoS pathogens 10(10): e1004423.

Tluckova K, Marusic M, Tothova P, Bauer L, Sket P, Plavec J, Viglasky V. 2013. Human papillomavirus G-quadruplexes. Biochemistry 52(41): 7207–7216.

Tosoni E, Frasson I, Scalabrin M, Perrone R, Butovskaya E, Nadai M, Palu G, Fabris D, Richter SN. 2015. Nucleolin stabilizes G-quadruplex structures folded by the LTR promoter and silences HIV-1 viral transcription. Nucleic acids research 43(18): 8884–8897.

van der Walt S, Colbert SC, Varoquaux G. 2011. The NumPy Array: A Structure for Efficient Numerical Computation. Comput Sci Eng 13(2): 22–30.

Wang SR, Min YQ, Wang JQ, Liu CX, Fu BS, Wu F, Wu LY, Qiao ZX, Song YY, Xu GH et al. 2016a. A highly conserved G-rich consensus sequence in hepatitis C virus core gene represents a new anti-hepatitis C target. Science advances 2(4): e1501535.

Wang SR, Zhang QY, Wang JQ, Ge XY, Song YY, Wang YF, Li XD, Fu BS, Xu GH, Shu B et al. 2016b. Chemical Targeting of a G-Quadruplex RNA in the Ebola Virus L Gene. Cell chemical biology 23(9): 1113–1122.

Waterhouse AM, Procter JB, Martin DMA, Clamp M, Barton GJ. 2009. Jalview Version 2-a multiple sequence alignment editor and analysis workbench. Bioinformatics 25(9): 1189–1191.

Zheng S, Vuong BQ, Vaidyanathan B, Lin JY, Huang FT, Chaudhuri J. 2015. Non-coding RNA Generated following Lariat Debranching Mediates Targeting of AID to DNA. Cell 161(4): 762–773.

Zhou B, Liu C, Geng Y, Zhu G. 2015. Topology of a G-quadruplex DNA formed by C9orf72 hexanucleotide repeats associated with ALS and FTD. Scientific reports 5: 16673.

